# *All fingers are not the same*: Handling variable-length sequences in a discriminative setting using conformal multi-instance kernels

**DOI:** 10.1101/139618

**Authors:** Sarvesh Nikumbh, Peter Ebert, Nico Pfeifer

## Abstract

Most string kernels for comparison of genomic sequences are generally tied to using (absolute) positional information of the features in the individual sequences. This poses limitations when comparing variable-length sequences using such string kernels. For example, profiling chromatin interactions by 3C-based experiments results in variable-length genomic sequences (restriction fragments). Here, exact position-wise occurrence of signals in sequences may not be as important as in the scenario of analysis of the promoter sequences, that typically have a transcription start site as reference. Existing position-aware string kernels have been shown to be useful for the latter scenario.

In this work, we propose a novel approach for sequence comparison that enables larger positional freedom than most of the existing approaches, can identify a possibly dispersed set of features in comparing variable-length sequences, and can handle both the aforementioned scenarios. Our approach, *CoMIK*, identifies not just the features useful towards classification but also their locations in the variable-length sequences, as evidenced by the results of three binary classification experiments, aided by recently introduced visualization techniques. Furthermore, we show that we are able to efficiently retrieve and interpret the weight vector for the complex setting of multiple multi-instance kernels.

## 1 Introduction

In various studies since the elucidation of the human genome, many different definitions of promoters have been used in different studies. For example, Butler et al. defined a core promoter as a minimal stretch of contiguous DNA sequence (∼ 40 nucleotides (nts)) that contains the transcription start site (TSS) and is sufficient for accurate transcription initiation [4], and a proximal promoter as a region in the immediate vicinity of the TSS, roughly 250 nts upstream and downstream. There are examples of many studies that consider either only an upstream region or using an arbitrary-sized window around the TSS (albeit fixed for the study) as promoter sequences. How does one know what size is appropriate in any independent new study or a study unifying such promoter sequences from multiple prior studies?

Discriminative machine learning methods like support vector machines (SVMs) [3] with their state-of-the-art performance on many relevant problems in computational biology (e.g., splice site prediction [15]) have been proven to be a very powerful tool. The earliest kernel-based approaches for computing similarities between biological sequences, e.g. spectrum [9] and mismatch kernel [10], allowed comparing sequences of different length, but they did not encode any positional information. Latter approaches, for example the weighted degree kernel [16] and oligo kernel [13], do consider positional information in the corresponding sequences, even with a certain amount of positional uncertainty [15]. Additionally, alignment-based sequence comparison also provides a position-dependent similarity score albeit with a gap penalty [17]. Thus, these approaches do allow deviations from exact matches but they are penalized. The oligomer distance histograms (ODH) kernel [11] allows comparing of sequences of different length by way of representing a sequence with a fixed-length feature vector. But it ignores information about the position of such oligomer pairs within the sequence.

These scenarios are outlined in Figure 1, panel Motivation, when comparing two sequences S1 and S2. Any position-aware kernel that also allows shifts can detect the signal in case (a) but not in case (b) where the signal is very far apart. Even if it does, it would penalize this deviation. Case (c) represents how ODH would detect this signal and thus consider the two sequences to be similar, but information on the position of this signal in the individual sequences is lost. This work is a step in the direction to tackle this issue: compare sequences allowing reasonable degree of positional freedom and not simultaneously penalizing this deviation or keeping it problem-dependent. This scenario can arise in case of Hi-C data where the pairs of loci interacting over a long-range are variable-length restriction fragments reported from the experiments and the causal signal in the two loci compared, for example, the enhancer and the promoter, does not have any positional restriction unlike the transcription start site in the promoter sequences.

In this work we approach the problem of handling variable-length sequences and allowing positional freedom when comparing them for problems such as identifying promoter architectures or analysis of long-range interaction partners collected from Hi-C experiments. We do so by breaking the individual sequences into segments (see Figure 1, panel Motivation, case (d)) and casting the problem into a multiple instance learning problem [6] where the instances in each bag are *parts* of the *whole* sequence. We employ conformal multi-instance kernels [2] to obtain the weightings for instances in each bag, thus rendering the capability to identify segments of a sequence informative for the prediction problem. We efficiently retrieve the weight vector for the complex setting of multiple conformed multi-instance kernels (outlined in section 2.3.1). We also demonstrate how to interpret the nonlinear classifiers by adopting visualization techniques that were recently introduced [14] in the more basic setting (outlined in section 2.3.2).

**Figure 1:**
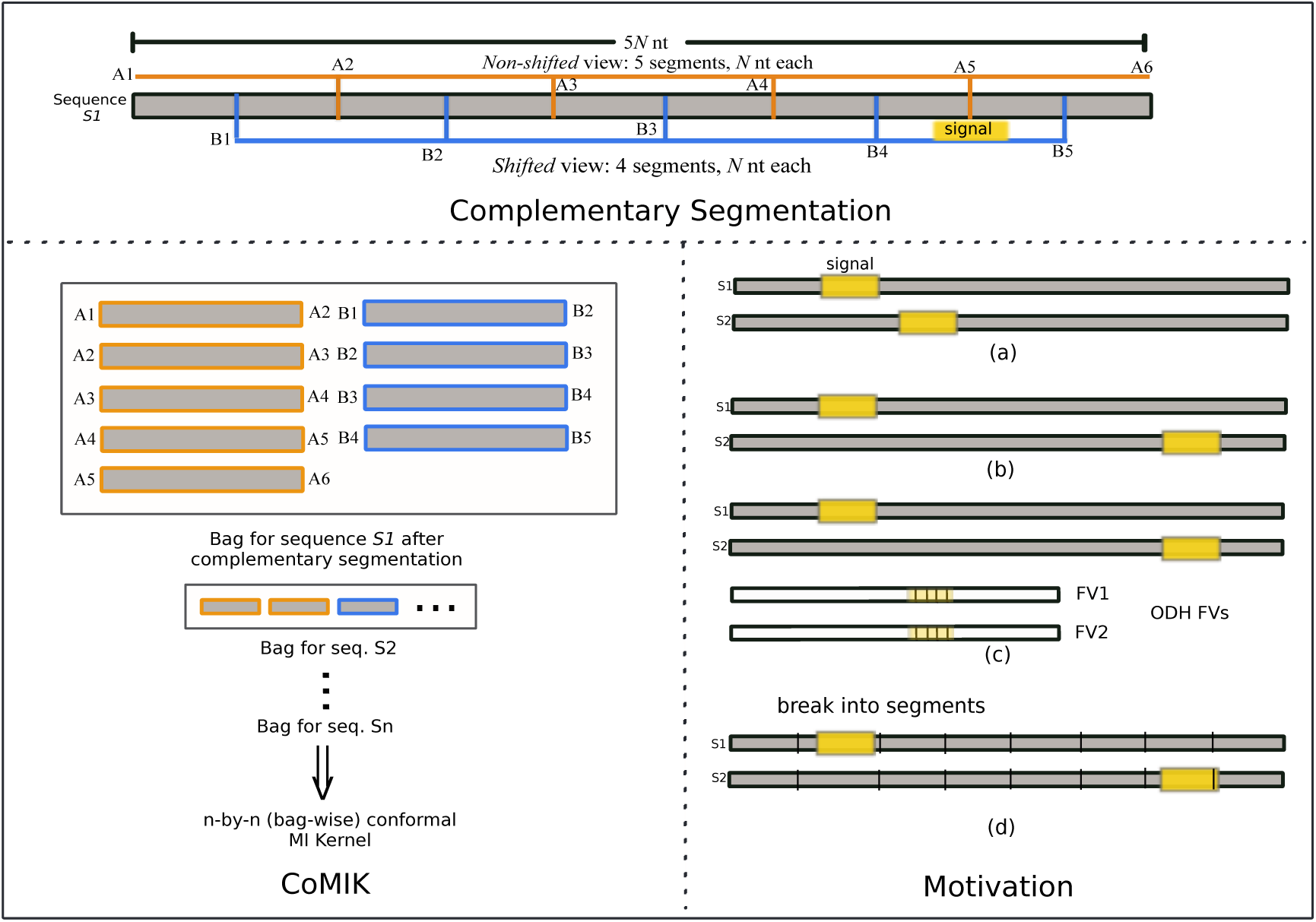
A schematic of our approach *CoMIK*, and the motivation. The complementary segmentation procedure is illustrated in the top panel. The sequence is shown in gray. *Non-shifted* segments are shown with an orange border, and *shifted* segments with blue border. The ‘Motivation’ panel shows the various scenarios when comparing sequences S1 and S2. Here, a signal in the sequence is shown as a yellow box. ‘FV1’ and ‘FV2’ denote the ODH (oligomer distance histograms [11]) feature vectors for S1 and S2 respectively. The panel ‘CoMIK’ shows a schematic of our approach starting from a sequence to the complementary conformal multi-instance (MI) kernel (see Methods for details). S1, S2, …, Sn are the *n* sequences in the collection.

## 2 Methods

Towards pair-wise comparison of variable-length sequences allowing positional freedom, we segment the individual sequences thus representing each sequence as a collection of its segments and then compare all segments of one sequence to all those of the other. With this, the typical binary classification problem involving sequences is cast into a multiple instance learning problem [6]. We call our approach *CoMIK* for ‘Conformal Multi-Instance Kernels’.

### 2.1 Segment Instantiation with Complementary Views

**Non-shifted Segment Instantiation:** Given any arbitrary length sequence, we propose representing it by its segments where a segment is defined as a smaller part of the whole sequence. Beginning right at the start of the sequence, we create segments of a predetermined size, along the sequence until it ends. The last segment is allowed to have a different size (either larger or smaller than other segments) to accommodate any remainder portion in case the sequence length is not an exact multiple of the segment size. We call this instantiation the *non-shifted* segment instantiation. A simple case of *non-shifted* segmentation is illustrated in Figure 1 (panel ‘Complementary Segmentation’). This segmentation provides the non-shifted view of the whole sequence as the first segment starts at the beginning of the sequence and, in total, the segments span the entire sequence.

**Shifted Segment Instantiation:** There may still be signals at the boundaries of any two non-shifted segments (see Figure 1, panel ‘Complementary Segmentation’, signal at position A5) which may get overlooked when comparing sequences using just *non-shifted* segments. To cover for this scenario, we introduce an alternate instantiation called *shifted* segmentation whereby the boundaries due to initial segmentation of the sequence end up in the same segment in this representation. In this case, segmentation begins from the mid-point of the first non-shifted segment, and proceeds to create further segments along the sequence essentially covering the boundaries of the non-shifted segments. The portions of the sequence before B1 and beyond B5 can be omitted since they are already covered in the *non-shifted* view (see Figure 1). *Shifted* segments can either be of same size as the *non-shifted* segments or different. Thus, *shifted* segmentation provides a complementary view of the same sequence covering the portions which get overlooked by *non-shifted* segmentation.

### 2.2 Complementary Set of Conformal Multi-Instance Kernels

Once segmented, we cast this problem into a multiple instance learning problem [6]. In the multiple instance learning setting, each sequence is thus treated as a bag and its segments as instances in the bag. One or more instances from a bag could be responsible for the bag to be classified as positive or negative due to the presence or absence of class-specific features. Since there is no restriction on the number of instances a bag can contain, this setting can inherently allow for considering arbitrary length sequences that result in an arbitrary number of instances per bag upon segmentation. Thus, there is a bag for each sequence in the collection containing *non-shifted* and *shifted* segments of the sequence.

#### 2.2.1 Multi-Instance Kernels

Gärtner et al. proposed the normalized set kernel (also known as the multi-instance kernel) for the multiple instance problem [8]. For each sample represented as a bag of instances, the kernel value between any two bags *X* and *X*′, *k*(*X*, *X*′), is given as in Eq. 1.

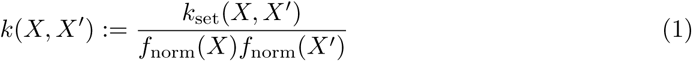

Here 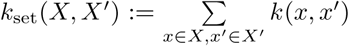 and *f*_norm_(*X*) is a suitable normalization function. One could normalize using either averaging (*f*_norm_(*X*) := #*X*, where #*X* denotes the number of instances in bag *X*) or feature space normalization 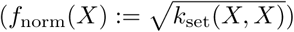. In this work, we used feature space normalization.

While the multi-instance kernel can successfully handle comparison between bags by comparing their individual instances, it has the issue that, in averaging, it looses any information related to the contributions of the individual instances. In other words, it treats all the instances in a bag equally. And, it is usually desirable to not only obtain a solution to a problem, but also to identify (a) the features that contribute to that specific solution, and (b) the parts which contain these features. Here, (b) amounts to knowing which instance(s) in a bag have features that helped determining the correct class label of the bag (positive or negative class). To this end, we propose using conformal multi-instance kernels [2] that allow us to obtain an instance weighting based on the contribution of these instances to learning the discriminant function.

#### 2.2.2 Conformal Multi-Instance Kernels

Blaschko and Hofmann proposed the conformal multi-instance kernel as a modification to the normalized set kernel [2]. This modification is a conformal transformation parameterized by *θ*, *t_θ_* > 0, applied to the kernel function, meaning that the transformation preserves the angle between vectors in the mapped space. The idea is to magnify those regions in the feature space which are discriminative and shrinking those which are not discriminative. Selection of these candidate regions in the feature space is done by clustering the complete set of input instances and choosing the corresponding cluster centres as candidate regions or expansion points. The decision of whether the region characterized by any cluster centre is discriminative or not is made by solving the multiple kernel learning problem as explained further.

Blaschko and Hofmann [2] proposed (a) the conformal transformation *t*_*θ*_(*x*) to be of the form given in Eq. 2.

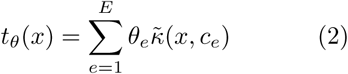

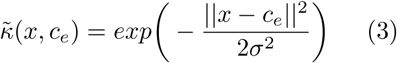

Here, *c_e_*’s denote the cluster centres indexed by *e* ∈ {1, …, *E*} for a total of *E* expansion points; and (b) 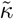 to be a Gaussian (Eq. 3) whose bandwidth (*σ*) can be adjusted. A large value of *θ_e_* in Eq. 2 denotes that the neighborhood of the corresponding expansion point *c_e_* is a discriminative region.

Thus, replacing *k*(*x, x′*) by its conformal transformation *t*_*θ*_(*x*)*t*_*θ*_(*x*′)*k*(*x*,*x*′)

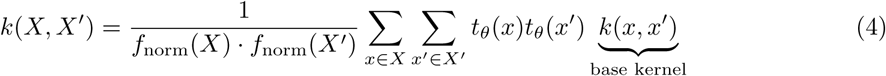

**Identifying expansion points.** To identify a set of expansion points, one could use *k*-means clustering with all available instances to identify *E*(=*k*) clusters, whose cluster centres, *c_e_* ’s, are then treated as expansion points. When dealing with too many instances, which could make the clustering process a bottleneck, Blaschko and Hofmann [2] suggest using the buckshot clustering approach [5] wherein, in order to identify *E* clusters from *n* instances, instead of using all *n* instances, one could perform *k*-means clustering using randomly sampled 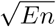 instances out of *n*. This has been shown to identify qualitatively similar clusters and being highly scalable at the same time [2].

**Resultant conformal multi-instance kernel.** Upon substituting Eq. 2 in Eq. 4, and simplification (see [2] for more details), the conformal multi-instance kernel is given by

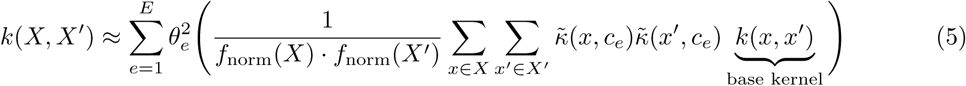

Eq. 5 is then posed as a multiple kernel learning (MKL) [1] problem (linear in 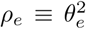) to simultaneously learn the SVM parameters *α* and *θ*.

**Obtaining individual instance weights.** Upon solving the MKL problem, once the sub-kernel weights (*θ_e_*’s) are obtained we can directly obtain *t_θ_*(*x*) for any instance *x* of a bag *X* using Eq. 2.

#### 2.2.3 Oligomer Distance Histograms (ODH) Kernel as Base Kernel

The choice of the base kernel to compare the individual instances depends on the problem. Here, we propose representing the individual segments of any sequence by its ODH representation [11] and using the ODH kernel [11] to compute similarities between them. In the ODH representation, any arbitrary-length sequence is represented by a feature vector that counts the occurrences of all pairs of short *K*-mers separated by *d* positions in the sequence.

For the DNA alphabet, 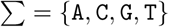, with 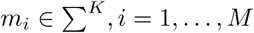 as all possible *K*-mers, any *L*-length sequence s can have *K*-mers separated by a maximum distance *D* = *L − K*. Thus, *d* − {0, …, *D*}. Here, distance between a *K*-mer pair is the difference between its starting positions in the sequence. For any *K*-mer pair (*m_i_, m_j_*), the distance histogram vector of its occurrences in sequence s is given as 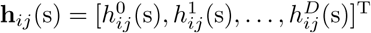. Here, each 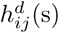 is the count of the number of times the *K*-mer pair (*m_i_, m_j_*) is observed at distance *d* in s. Finally, the feature space transformation of sequence s is obtained by stacking together the distance histograms of all *K*-mer pairs over Σ.

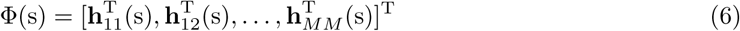

Then, *N* training samples are given as: 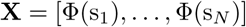 and the *N × N* kernel matrix is given by **K** = **X**^T^ **X**.

### 2.3 Interpretation and Visualization of Features

In the following, we discuss how one can interpret and visualize the sequence features deemed important by *CoMIK* for a prediction problem.

#### 2.3.1 Obtaining the SVM Weight Vector for *CoMIK*

In the MKL problem [1], the weight vector corresponding to a given sub-kernel *Km* is given as in Eq. 7.

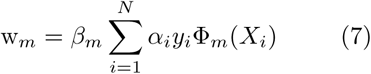

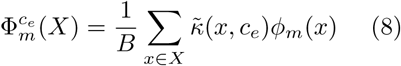

Here *β_m_* is the sub-kernel weight learnt by solving the MKL problem and each Φ*_m_* (*X_i_*) is the feature space representation of sequence *X_i_* corresponding to sub-kernel *K_m_*. And, for the conformally transformed multi-instance setting, this would mean Φ*_m_* (*X*) is the bag-level, transformed ODH representation of the sequence corresponding to the cluster centre chosen when computing the kernel *K_m_*. Thus, 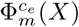 can be represented mathematically as in Eq. 8 where *ϕ_m_*(*x*) is the ODH representation of segment *x* (Eq. 6) belonging to bag 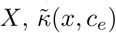 is the Gaussian transformation (Eq. 3) and *B* is the feature space normalization factor. Following [20], *B* can either be 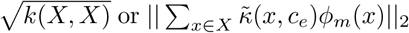 since our base kernel, the ODH kernel, is a dot product kernel (refer to section 2.2.3). Thus, we have a bag-level representation of a sequence corresponding to all cluster centres which allows us to compute all the relevant weight vectors. These individual weight vectors can also be used to make fast predictions on test examples. For this, we only need the transformed ODH representations of the test examples corresponding to each kernel in the collection.

#### 2.3.2 Visualizing Features from the *CoMIK* Weight Vector

Figure 2 shows two ways of visualizing the features deemed important by *CoMIK* in discerning the positive set of sequences from the negative set. The bottom-left panel in Figure 2 shows the distance-centric view, the ‘Absolute Max Per Distance’ (AMPD) visualization [14], and, the right panel, *K*-mer-centric view [11]. In the AMPD visualization, based on the ODH feature representation, each dimension of the weight vector corresponds to the histogram count of an oligomer pair lying at a given distance *d* (unit: basepairs) (refer section 2.2.3, Eq. 6). From this weight vector, for each distance value, among all the *K*-mer pairs, we pick the pair that is assigned the most positive and most negative coefficient. A positive coefficient value means the feature (i.e., the *d*-separated *K*-mer pair, *d* ∈ {0, 1, …, *D*}) is prominent among the positive set of examples, otherwise negative. This provides a distance-centric view of the important features. The *K*-mer-centric view [11] shows the role of each *K*-mer pair towards prediction. Simply stated, the *K*-mer-centric view of the discriminant is a matrix which is obtained upon performing, for all *K*-mer pairs, an *ℓ*_2_-norm of the relevant dimensions of the weight vector, corresponding to all distance values considered, with itself. Thus, a pair which holds high importance (i.e., it has large coefficients in the discriminant, positive or negative) will have higher absolute value in the matrix.

## 3 Materials

**Simulated data set:** We prepared a simulated data set with sequences of arbitrary length and with a mix of many coupled and non-coupled motifs. These are shown in Table 1. Three sets of motifs were planted in the sequences in this data set. In each motif set, the serial numbers marked with `P` and `N` (e.g., `4P` and `4N` in set A) denote the variants planted only in the positive and only in the negative examples, respectively. The differences in these variants are underlined.

**Table 1:**
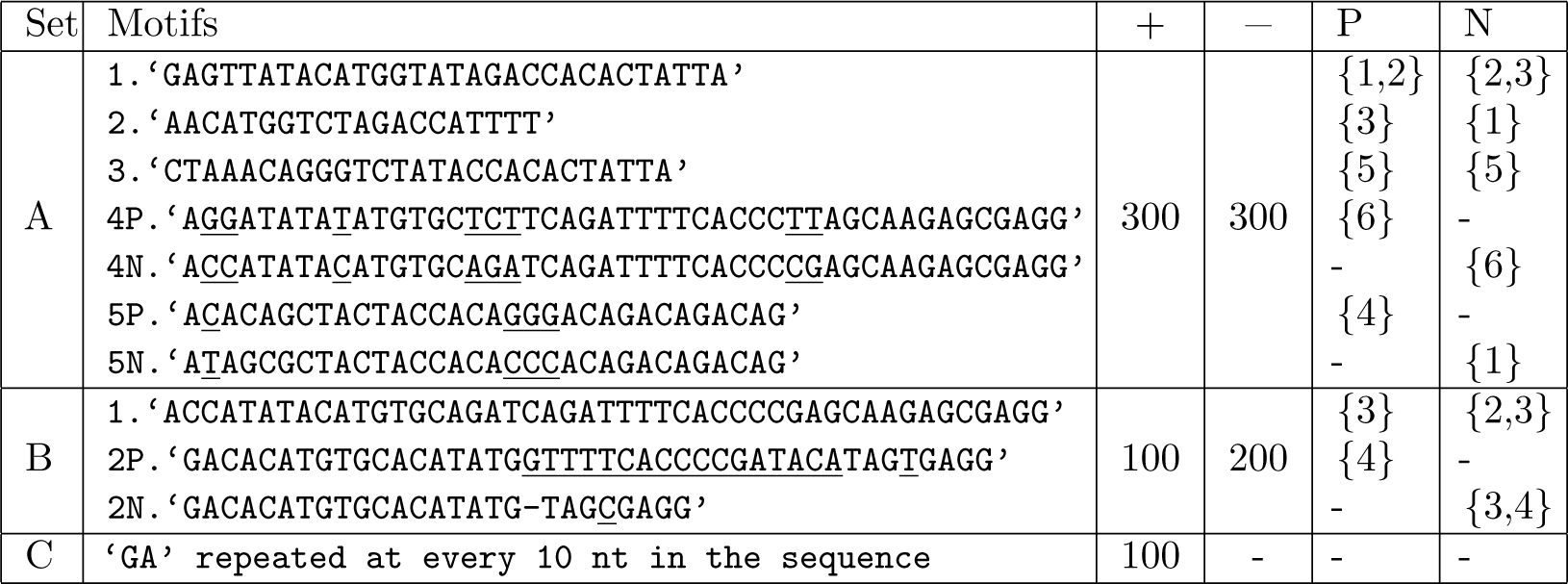
Motif sets planted in the simulated data set. Columns marked ‘+’ and ‘−’ give the number of positives and negatives respectively containing the corresponding set of motifs. Columns ‘P’ and ‘N’ give the #segment (non-shifted) in which the motif could lay (start positions).

The motif `2N` in motif set B has a gap compared to `2P`. This gap is denoted by a ‘-’. All other motifs in sets A and B (motifs 1, 2 and 3 in set A, and motif 1 in set B) are present in both positive and negative examples. The occurrence positions of all motifs in either the positive or negative set could differ since the sequences are of varying length. Based on the windows inside which the various motifs could lay in the sequence, the corresponding 70nt-segment numbers (for non-shifted segments) are given in Table 1. Motif set C is motivated from the case of Drosophila GAGA factor association to closely spaced (GA)*n* elements *in vivo* [21]. The positive and negative set each have 500 sequences. The different kinds of positive and negative sequences have different length distributions: in the range [300,500]nt for sequences of types A and B, and [500,600]nt for type C. All sequences are generated such that nucleotides `A, C, G` and `T` have a uniform distribution and the planted motifs have 0.1 mutation probability. Maintaining equal proportions of the different kinds of positives and negatives, we held out 200 sequences as unseen test examples (100 positives and 100 negatives) and used the remaining 800 sequences for training.

**Yeast:** Lubliner et al. studied yeast core promoter sequences analyzing the effect of sequence variation in different core promoter regions [12]. Among other things, the authors showed that location, orientation, and flanking bases are important for TATA element function. We obtained a total of 316 118nt-long core promoter sequences ([-118,-1] relative to the TSS) for which the core promoter activity measurements were provided and followed the procedure in Figure 5 in [12] to classify them into two classes– sequences showing either low or high activity (low or high expression), giving 28 positive and 288 negative sequences.

**5C:** In a recent study, Nikumbh and Pfeifer [14] approached the problem of predicting the long-range interaction partners of a genomic locus (of interest) profiled in 5C experiments in cell lines GM12878, K562 and HeLa-S3 [18] using the DNA sequence at the interacting and non-interacting loci. For any 5C-profiled TSS-containing region, the distal loci that showed a significant interaction with it in all replicates were considered as positive and the ones that did not interact significantly in any replicate were considered negative. We fetched the positive and negative set of sequences for one region (*region* 0) per cell line [14].

## 4 Experimental Setup

For each data set, we performed 5-fold nested cross-validation (CV) by splitting the data into 80%:20% for training and test respectively. For each outer-fold, model selection was performed with a 5-fold inner CV loop on the training set with *ℓ*_1_ - and *ℓ*_2_-norm MKL. We note that *CoMIK* accounts for any class imbalance by proportionately up-weighting the misclassification cost for the minority class as proposed in [7]. All parameters and the range of values tested for them are given in Table 2. Of these, #Clusters, *σ* and SVM-cost are optimized by cross-validation while other parameters, namely segment size, oligomer length and maximum distance, are assigned fixed values for each individual run. We used the same segment-size for the *shifted* and the *non-shifted* cases. The best performing set of parameter values obtained from the inner CV-folds was used to re-train the model using the complete training data and make predictions on the unseen test set of examples per outer CV-fold. We report the area under the receiver operating characteristic (ROC) curve (AUC) for predictions on this held-out test set averaged over the five outer folds.

**Table 2:** Parameters and the range of values tested for the simulated, 5C and the yeast data set.

| Parameters | Simulated data set | 5C | Yeast |
| --- | --- | --- | --- |
| #Clusters | {2, 5, 7} | {5, 7, 10} | {2, 5, 7} |
| Segment size | {50, 70} | 50 | 10 |
| Sigma ($\sigma$) for Gaussian transformation | $10^{\{-1, \dots, 2\}}$ | $10^{\{2, 4, 6\}}$ | $10^{\{-1, \dots, 2\}}$ |
| Oligomer length | {2, 3} | {2, 3} | {2, 3} |
| Maximum distance | {50, 70} | 50 | 10 |
| SVM-cost | $10^{\{-3, \dots, 3\}}$ | $10^{\{-3, \dots, 6\}}$ | $10^{\{-3, \dots, 3\}}$ |

We compare our performance on the simulated data set to that of KIRMES (Kernel-based Identification of Regulatory Modules in Euchromatic Sequences) [19]. KIRMES was shown to perform well on gene sets derived from microarray experiments for identifying loss or gain of gene function [19]. For a collection of positive and negative genomic sequences, given a set of motifs representing transcription factor binding sites and their match-positions (obtained by performing a motif-finding step a priori) in the sequences, KIRMES picks a *fixed-size* window around the best match-position of the motif in the sequence as a representative of the sequence for that motif. These selected, *fixed-size* portions from all sequences (thus, equal length) are used to compute a WDSC kernel (weighted degree kernel with shifts and conservation information; see [19] for complete details) corresponding to that motif. This procedure results in as many kernels as the number of motifs. The remaining parts of the sequences (those not selected for any motif) are *neglected*.

## 5 Results

In the following, with computational experiments on a simulated data set, and yeast and 5C data, we demonstrate how *CoMIK* can be used to uncover the features regarded important for classification together with their locations (at the segment level) in any candidate sequence.

**Simulated data set:** For this data set, while KIRMES achieves an AUC of 0.9432, *CoMIK* attains near-perfect classification, AUC 0.9960 *±* 0.003. We surmise that the superior performance of *CoMIK* is due to the sequences containing the dinucleotide repeat motif ‘`GA`’ (see Figure 1) which may not be captured at the motif-finding step and thus affect KIRMES’ prediction.

We provide visualizations from the run that achieved the best performance with oligomer-length 3, segment-size 70nt, *ℓ*_1_-norm MKL in Figure 2. The top panel visualizes the 70nt-long segments of 50 out of the 200 test sequences horizontally. For each sequence, the *non-shifted* segments are followed by its shifted segments. Per sequence, the higher-ranked segments would be the ones where the features are located. Figure 2, bottom-left panel, is a distance-centric visualization of the SVM weight vector and the bottom-right panel, the *K*-mer-centric view. While the K-mer-centric view clearly indicates `GA`'s important role, the distance-centric visualization shows that it could be periodic. Experiments using different segment-sizes can easily uncover the fact that they are spread throughout the sequences.

**Figure 2:**
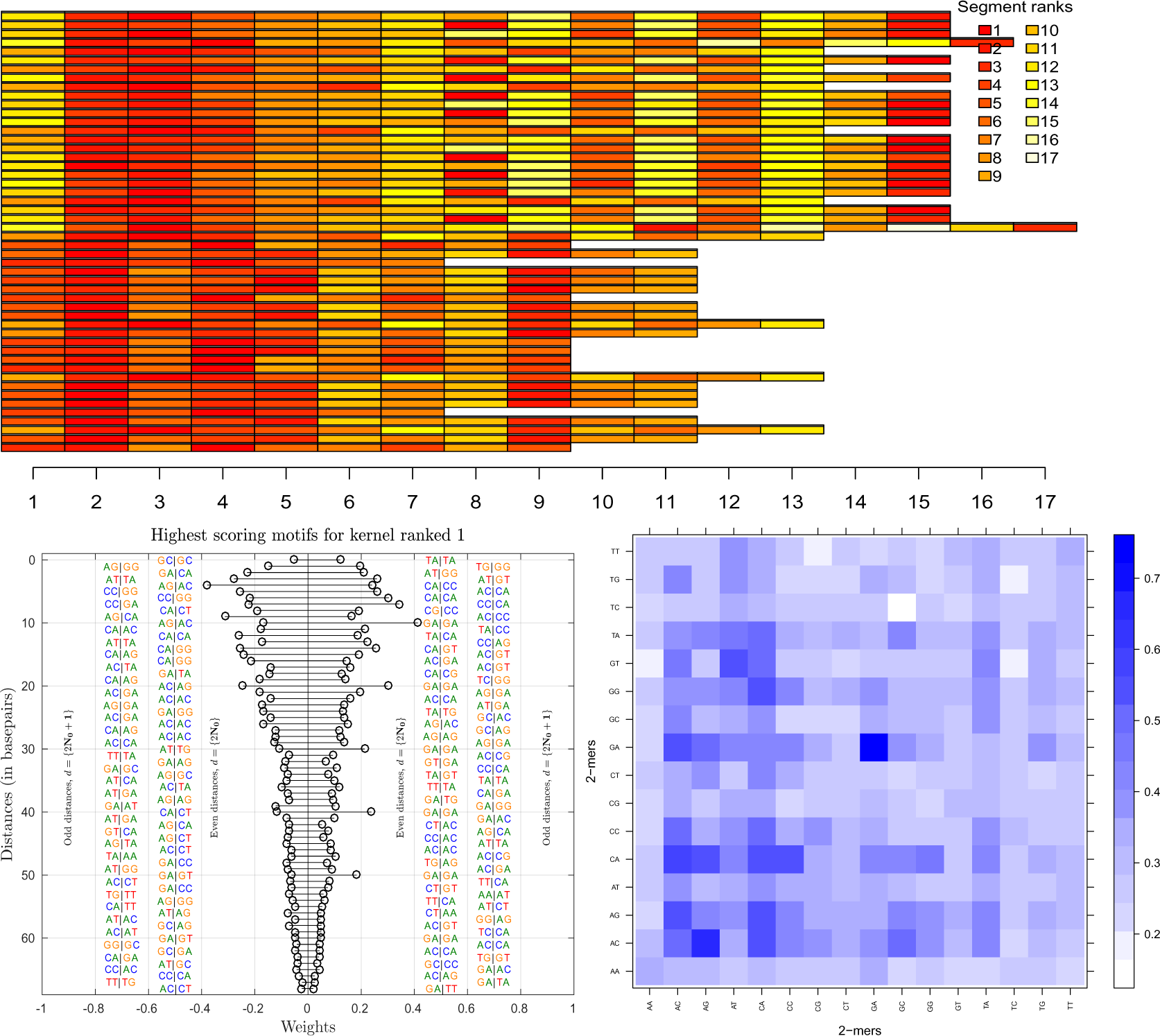
Visualizing the weights assigned to segments of the variable-length sequences in simulated data set, distance-centric (bottom-left panel) and *K*-mer-centric visualizations (bottom-right panel) of features for the simulated data set.

**Yeast:** *CoMIK* achieved an AUC of 0.9459 ± 0.029 on this data set. Furthermore, the most important features represent motifs known as important for classification. From the run that achieved best performance with segment-size 10nt, oligomer-length 3, 7 clusters and 1\-norm MKL, we visualized the 3-mer pairs deemed important by *CoMIK* for this classification in Figure 3, left panel. The right panel here visualizes the sequences and their segments as a heatmap. The 316 sequences are arranged vertically from top to bottom, and their segments horizontally. For the 118nt-long sequences in this data set, the segment-size of 10nt lead to 12 *non-shifted* and 11 *shifted* segments, and are arranged in that order. Thus, the coordinates for the *non-shifted* and *shifted* segments in the sequence are as marked on the top of the heatmap. We observe that segments 3 and 9, i.e., regions [-98,-89] and [-38,-29] happen to be ranked first consistently. Segments 15 and 21 are the best-ranked shifted segments also corresponding to the same genomic window. And, indeed, Lubliner et al. report that the main TSS lay at position -30 and that the regions [-118,-99] and [-98,-69] hold important features which upon mutations greatly reduced expression [12]. In the left panel, the top-ranked kernel shows TATA-like elements to be important for classification. Furthermore, among the features reported by other kernels in the collection (not shown), *CoMIK* rightly identifies `T/C`-rich K-mers to be enriched among the positive sequences as against `G/C`-rich K-mers which are also reported in Supplementary Figure 4 in [12].

**5C data set:** Performances of *CoMIK* on the three cell lines are given in Table 3. For comparison with the method by Nikumbh and Pfeifer [14] that represen ts the complet e restriction fragment with its ODH representation, we directly report the performances with oligomer length 3 from Table 1 in [14]. We observe that in experiments with segment-size 50 using 3-mers, *CoMIK* already achieves as good or better performance. Furthermore, the additional ability of *CoMIK* to identify important portions in the individual sequences could give novel insights.

**Figure 3:**
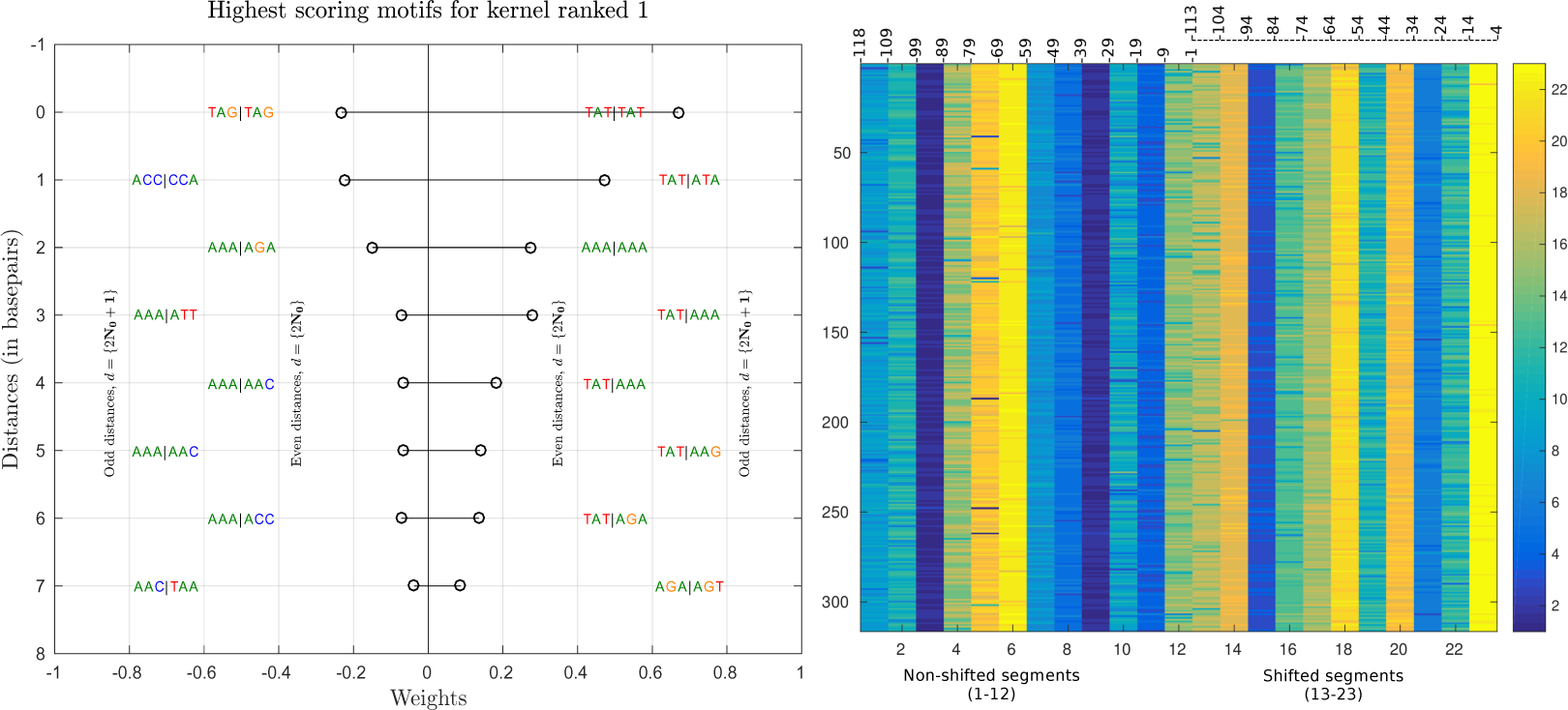
Distance-centric visualization of features (left) and visualization of weights assigned to segments per sequence for the yeast data set.

**Table 3:** 5C data set results: Test AUC values (mean±s.d.) for *region* 0 [14] in three cell lines.

| Method/Cell lines→ | GM12878 | K562 | HeLa-S3 |
| --- | --- | --- | --- |
| Nikumbh and Pfeifer [14] | 0.7417 ± 0.059 | 0.8163 ± 0.071 | 0.6914 ± 0.058 |
| *CoMIK* | 0.7829 ± 0.063 | 0.7920 ± 0.084 | 0.6993 ± 0.012 |

## 6 Discussion

We presented a multiple instance learning-based approach, called *CoMIK* (‘Conformal Multi-Instance Kernels’), that can handle highly dispersed features in comparing variable-length sequences in a discriminative setting. We assessed the performance of *CoMIK* on three classification problems: a simulated data set and t wo real biological data sets including a 5C data set. Together with the visualizations, we demonstrated the efficacy of *CoMIK* in all these problems. As compared to KIRMES, where the classifier completely relies on the motif-finding step *apriori* for its input, *CoMIK,* by design, uses the complete sequence and is able to locate the portions deemed important for the prediction problem. This enables *CoMIK* to avert the risk of ignoring the lo w-affinity, weak binding sites in the sequences which can be missed by KIRMES. Technically, one could use the complete sequence with KIRMES provided the set of motifs considered are spread through-out the sequence, but that is again controlled by the motif-finding stage. *CoMIK* allows positional freedom in comparing sequences. For the 5C data set, Nikumbh and Pfeifer also used the ODH representation to compare the restriction fragments [14], but their approach does not give any information on the location of the features in the long restriction fragments. Our results on this data set showed that in this scenario looking closely at shorter segments rather than the complete restriction fragments can help attain better performance. Additionally, *CoMIK's* ability to locate signal within the sequence could be useful in studying the so-called structural interactions between the intervening chromatin [18] of the long-range interacting loci.

We note that *CoMIK's* computation time is largely governed by the clustering step and the subsequent transformation of the segments - both performed at every CV iteration. Our implementation exploits the sparsity of short individual segments, makes use of sparse representations and com putations. In the clustering step, the buckshot heuristic is incognizant of the imbalance prevalent in the data. This could be improved by using a stratified sample with buckshot clustering. We also note that for scenarios wherein positional information is important, kernels like the WDS [15] or the oligo kernel [13] could be more suitable as base kernels depending on the problem.

